# QTL mapping of oat crown rust resistance in Australian fields and identification of a seedling resistance locus in oat line GS7

**DOI:** 10.1101/2025.08.24.671654

**Authors:** Duong T. Nguyen, David Lewis, Eva C. Henningsen, Zhouyang Su, Rohit Mago, Jana Sperschneider, Peter N. Dodds, Allan Rattey, Belayneh A. Yimer, Kathy Esvelt Klos, Melania Figueroa

## Abstract

The development of oat cultivars with resistance to crown rust caused by *Puccinia coronata* f. sp. A*venae* (*Pca*) is key for sustainable disease control. This study examined two recombinant inbred line populations, Provena x GS7 and Boyer x GS7, to identify adult plant resistance QTL in Australian fields. Seven distinct QTL associated with rust resistance were identified, with KASP markers developed for single nucleotide polymorphisms (SNPs) tightly linked to the four most significant QTL on chromosomes 4A and 7A. A major QTL named *QPc_GS7_4A.2* with a resistance allele derived from line GS7 was mapped to chromosome 4A, overlapping with genomic regions previously associated with both resistance gene *Pc61* and adult plant resistance. Genetic mapping for rust resistance at seedling stage using a subset of Provena x GS7 lines with contrasting alleles at *QPc_GS7_4A.2* confirmed the role of this locus on seedling resistance, likely by *Pc61*. Furthermore, we found similar resistance profiles between GS7 and the Pc61 differential line against 20 *Pca* isolates at the seedling stage. Haplotype analysis of *QPc_GS7_4A.2* in the oat crown rust differential set and an oat collection revealed the resistance haplotype in lines previously postulated to carry resistance gene *Pc61*. These results suggest that the QTL *QPc_GS7_4A.2* is closely linked to the *Pc61* locus on chromosome 4A. The KASP markers associated with *Pc61* and QTL identified in this study will be valuable tools, allowing breeders to efficiently integrate the resistance allele for gene combinations in new cultivars, particularly in regions where *Pc61* remains effective.

## Introduction

Crown rust, a fungal disease caused by *Puccinia coronata* f. sp. *avenae* (*Pca*) is a significant threat to oat production worldwide (Nazareno et al. 2018; Simons 1985). Major resistance (*R*) genes conferring race-specific resistance against *Pca* have been widely used in oat breeding to safeguard crops from this disease. Such *R* genes usually encode specific receptors in the host that recognise effector molecules, which can vary between pathogen races, explaining the race-specificity (Dodds et al. 2024; Jones et al. 2024). This type of rust resistance is usually expressed from the seedling stage and lasts throughout the plant life cycle, so it is called all-stage resistance (ASR) (Periyannan et al. 2017). However, the effectiveness of ASR genes often diminishes over time due to the emergence of variants in the pathogen population to overcome resistance (Figueroa et al. 2020). Oat cultivars with ASR genes for crown rust typically lose resistance within five years of release, a pattern likely to persist with continued reliance on race-specific resistance genes (Carson 2008). To combat this challenge, researchers and breeders have switched their focus to identifying and harnessing sources of more durable resistance.

Another class of rust *R* genes defined in cereals confer non-race specific resistance associated with a partial resistance phenotype. Partial resistance is considered highly effective in managing the disease because it slows down the evolution of pathogen virulence. Unlike race-specific ASR, partial resistance is not limited to specific pathogen races and operates at the adult plant stages, therefore it is called adult plant resistance (APR) (Ellis et al. 2014; Periyannan et al. 2017). While APR does not entirely inhibit fungal sporulation, it curtails pustule size, and spore production, and prolongs the latent period (Portyanko et al. 2005). However, incorporating APR into breeding programs presents challenges due to its quantitative nature, and multiple loci must be combined to achieve high levels of resistance (Nazareno et al. 2022). Therefore, introgression of these novel alleles into elite germplasm is a lengthy process. Nevertheless, several major loci governing APR have been identified offering potential shortcuts in breeding for durable disease resistance. For instance, the genes *Lr34* and *Lr67* in wheat encode membrane transporters that suppress rust growth independently of specific recognition (Krattinger et al. 2009, 2016; Milne et al. 2019; Moore et al. 2015).

In oat, of approximately 100 loci conferring resistance to *Pca* that have been catalogued to date, six are associated with APR (*Pc27*, *Pc28*, *Pc69*, *Pc72*, *Pc73*, *Pc74*). However, the chromosomal locations for all these genes are unknown (Carson 2017; Harder et al. 1984). Additional sources of APR have been postulated in various cultivars (Cabral et al. 2011; Heagle and Moore 1970; Luke et al. 1972; Upadhyaya and Baker 1962; Welsh et al. 1953). Although most of these APR sources remain uncharacterised, and their underlying genetic mechanisms are unknown. So far, few molecular markers associated with APR in oat have been developed for use in breeding and selection (Lin et al. 2014; Nazareno et al. 2022; Rines et al. 2018).

Previously, Babiker et al. (2015) identified three APR QTL in three mapping populations of recombinant inbred lines (RILs) derived from crossing the partially resistant sources, CDC Boyer (referred to as Boyer hereafter) and a breeding line 94197A1-9-2-2-2-5 (referred to as GS7 hereafter) with the susceptible cultivar Provena, and with each other (Boyer x GS7). In their study, Babiker et al. (2015) found that Boyer contributed the resistance alleles of two QTL located on chromosomes 12D (intervals of approximately 15.8 cM) and 19A (9.7 cM) (based on the cytology-based nomenclature by Sanz et al. 2010), while GS7 contributed one QTL on chromosome 13A (15.4 cM). These correspond to positions on chromosomes 2Ds, 4As and 7As, respectively, under the uniform nomenclature system of Jellen et al. (2024).

This study aimed to evaluate the effectiveness of APR resistance loci from Boyer and GS7 under Australian field conditions using the same RIL populations. DArTSeq genotyping was employed to enhance marker density to resolve QTL regions. A total of seven distinct QTL associated with oat crown rust resistance were detected, and KASP markers were developed for single nucleotide polymorphisms (SNPs) tightly linked to the four most significant QTL on chromosomes 4A and 7A. KASP assays were implemented to assess the presence of the resistance alleles in an oat collection of 182 lines, including 150 lines from Nguyen et al. (2023) and 32 lines postulated to carry APR. A strong QTL from GS7 linked to crown rust resistance was identified on homoeologous regions of chromosomes 4A and 4D, overlapping with genomic regions previously linked to the all-stage resistance gene *Pc61* and adult plant resistance. Genetic mapping for rust resistance at the seedling stage using a subset of Provena x GS7 RILs with contrasting alleles at *QPc_GS7_4A.2* confirmed that this locus contains a seedling resistance gene. Haplotype analysis using SNPs linked to this QTL in the differential set (Henningsen et al. 2024) and designed KASP markers in the oat collection identified the *QPc_GS7_4A.2* resistance haplotype as highly specific to *Pc61* carriers. The results suggest a strong linkage or potential identity between the *QPc_GS7_4A.2* on chromosome 4A and the *Pc61* locus. Overall, these findings underscore the potential of GS7 and Boyer as a valuable sources of crown rust resistance, with the identified QTL and associated KASP markers providing insights to uncover underlying mechanisms and support marker-assisted selection in breeding programs.

## Materials and methods

### Plant material

The recombinant inbred lines (RILs) from the two mapping populations, ‘Provena x 94197A1-9-2-2-2-5’ and ‘CDC Boyer x 94197A1-9-2-2-2-5’, referred to as Provena x GS7 (n=91) and Boyer x GS7 (n=98), respectively, were developed and described by Babiker et al. (2015) through eight generations of selfing after F2 generation. These RILs were sourced from the USDA Agricultural Research Service, Aberdeen, ID, USA. This study also includes an oat collection of 182 lines, comprising 32 lines postulated to carry APR and 150 lines recently compiled by Nguyen et al. (2023). These lines were sourced from USDA-ARS (St. Paul, MN, USA), the Australian Grain Genebank (AGG), and the CSIRO Avena seed stock. The list of plant materials is included in **Supplementary File 1, Table S1 and Table S2**.

### Disease resistance phenotyping

Field infection data for Provena x GS7 and Boyer x GS7 RILs was collected from field trials in Australia at Manjimup, WA (33.24° S, 116.16° E), in 2023 and 2024, and Cobbitty, NSW (33.99° S, 150.69° E), in 2024. Data for the oat collection was recorded in Manjimup (2023) and Cobbitty (2024). In all trials, lines were planted 1-row wide x ∼0.5m long. In the Manjimup trials in 2023 and 2024 (MJ23 and MJ24), rust races (triplet codes as described by Park 2000) 0001-2 and 0005-0 were used, while in the Cobbitty trial 2024 (CB24), the pathotypes “4473-4,6,10, Bett, Barc”, “0767-3,4,5,6,10,Wa,Vo”, and “3707-1,4,5,6,7,10,12,Wa,Nu,Gw,Ge,Dr,Al” were utilised as they are highly frequently found on those regions. Around the flowering between Zadoks growth stage GS59 - GS80 (Zadoks et al. 1974), rust infection severity was scored using the 0-to-100 modified Cobb scale (Peterson et al. 1948) in Manjimup, whereas in Cobbitty, severity was rated on a 1–9 scale. To enable comparison between the two locations, the 1-9 scale scores from CB24 were converted into percent severities following the method of Bariana et al. (2007).

Rust infection assays at the seedling stage were conducted in growth cabinets on a subset of 30 RILs from the Provena x GS7 population, which were selected for having contrasting alleles at the QTL *QPc_GS7_4A.2* identified through field-based QTL mapping. These lines were tested against *Pca* isolate 22WA54, which was virulent Provena but avirulent to GS7 to evaluate the effect of the locus *QPc_GS7_4A.2* at the seedling stage. In addition, similar seedling assays were also performed on GS7, Provena, Pc60, Pc61, and Swan oat lines against 20 *Pca* isolates to compare their resistance profiles (Henningsen et al. 2024; Nguyen et al. 2025). Plants were grown under controlled conditions (23/18°C, 16/8hrs, light/dark) and *Pca* spore samples were applied to plants 10 days after sowing, and infection scores were recorded 10 days post-inoculation. Seedling infection and scoring methods were described by Miller et al. (2020), with infection type scores (“0”, “0;”, “;”, “;C”, “1;”, “1”, “2”, “3”, “3+”, “4”) converted to a 0-9 numeric scale (0, 1, 2, 3, 4, 5, 6, 7, 8, 9) respectively for plotting as heatmaps using ComplexHeatmap v2.20.0 in R v4.4.0 (Gu 2022).

### Genotyping

Five seeds from each oat line were sent to Diversity Arrays Technology Pty Ltd. (Canberra, Australia; https://www.diversityarrays.com/) for DNA extraction and genotyping using their proprietary genome complexity reduction-based sequencing technology. To identify SNP markers, the sequences obtained from DArTSeq were aligned against the reference genome sequence *Avena sativa* OT3098 v2 (PepsiCo, https://wheat.pw.usda.gov/jb?data=/ggds/oat-ot3098v2-pepsico), deposited in the GrainGenes database (Yao et al. 2022), using BLAST (Altschul et al. 1990) with an expected value (E) threshold of less than 5e-7 and sequence identity greater than 70%. Only markers that matched a genomic location were kept. The SNP dataset was filtered in dartR (v2.9.7) by first removing loci with all missing data, then excluding monomorphic loci to retain only polymorphic markers (Gruber et al. 2018). A call rate filter removed loci with over 50% missing data and markers with a minor allele frequency below 0.01 were discarded. The genotypic data of Provena x GS7 and Boyer x GS7 RILs were transformed to a parent-based format (ABH) by using the GenosToABH plugin from TASSEL (v.5.2.64), using the codes A: male parent, B: female parent, H: heterozygous. Data imputation was conducted using ABHgenotypeR v.1.0.1 R package (Furuta et al. 2017).

### Linkage map construction for Provena x GS7 and Boyer x GS7 RILs

Linkage group construction was carried out using the “mstmap” algorithm implemented in the R package ASMap v.1.0.7 R package (Taylor and Butler 2017). The initial group assignment was based on *p*-value thresholds of 1e-19 for Provena x GS7 and 1e-21 for Boyer x GS7, with the “Kosambi” function used for genetic distances. Markers not assigned to linkage groups were removed. Subsequently, chromosome names were assigned to linkage groups by identifying the most frequent chromosome location of DArTSeq markers mapped to the OT3098 v2 reference genome. Linkage groups with the same chromosome name were then merged using the “mergeCross” function with a gap threshold of 10 cM, and marker order was refined through a second round of “mstmap” using a p-value threshold of 1e-6. Linkage groups with zero length or fewer than seven markers were filtered out. Unmerged linkages were merged with other already-merged linkage groups sharing the same chromosome name using a permissive p-value threshold of 1e-2, considering orientation and correlation with the physical map. Genetic distances were recalculated using the “Kosambi” function after merging. The correlation between genetic map marker order and the reference genome was calculated using a Pearson’s correlation test and visualized with the “ggplot2” R package (Wickham 2016).

### QTL mapping

QTL mapping was performed for the Provena x GS7 and Boyer x GS7 RILs using the genetic maps and field phenotypic data. For the subset of RILs from Provena x GS7 (n=30), mapping was carried out using the genetic map and seedling resistance data. All analyses were conducted using composite interval mapping (CIM) with the R/qtl package (v1.66) (Haley-Knott with forward selection to three markers and a window size of 10 cM) (Broman and Sen 2009). The threshold for the logarithm of odds (LOD) for significant QTL declaration was defined by 1,000 permutations at *p*≤0.05. The custom interactive R script for QTL mapping, developed in RStudio 2024.04.2, is based on the R/qtl manual by Broman and Sen (2009). The percentage of genotypic variance explained (GVE) was estimated using the formula 1 – 10^−2 LOD/*n*^, where n is the sample size and LOD is the LOD score of the QTL (Broman et al. 2003).

### Comparative mapping analysis

The flanking sequences of significant markers associated with QTL from previous studies (Babiker et al. 2015; Klos et al. 2017; Nazareno et al. 2022; Rines et al. 2018; Wight et al. 2004) were searched against the reference genome OT3098 v2 by BLAST using Geneious Prime® 2022.2.2. The flanking sequences of markers were taken from NCBI GenBank (https://www.ncbi.nlm.nih.gov/nuccore/) and T3/Oat (https://oat.triticeaetoolbox.org/). Syntenic relationships between chr4A and 4D were established and visualised through a Comparative Genomics Platform (CoGe; https://genomevolution.org/coge/SynMap.pl) and the SynMap2 tool (Haug-Baltzell et al. 2017).

### KASP assays

We designed KASP markers for the SNPs closely linked to identified QTL: *SNP-350765064_4A*, *SNP-351486359_4A*, and *SNP-352361337_4A* for *QPc_GS7_4A.1, SNP-459779133_4A, SNP-459972095_4A,* and *SNP-460562184_4A* for *QPc_GS7_4A.2, SNP-4039736_7A* and *SNP-411798_7A* for *QPc_GS7_7A,* and *SNP-67318388_7A* and *SNP-69787185_7A* for *QPc_Provena_7A/QPc_Boyer_7A*. The flanking sequences of these SNPs were imported into the Kraken™ software system for marker design using the default parameters (LGC Biosearch Technologies, UK; https://www.biosearchtech.com/). KASP assays were performed in the SNPline PCR Genotyping System (LGC, Middlesex, United Kingdom), following the methods described by Shi et al. (2023). KASP primer sequences are included in **Supplementary File 1 Table S3.**

### Haplotype visualisation

Haplotype of QTL *QPc_GS7_4A.2* identified in chr4A from mapping analysis was visualised in the oat collection and the differential set using Flapjack (Milne et al. 2010). The *QPc_GS7_4A.2* haplotype in the oat collection is visualised based on the result of KASP genotyping assays of *KASP_4A_459779133,* KASP_4A_459972095, and KASP_4A_460562184; and the *QPc_GS7_4A.2* haplotype in the differential set (Henningsen et al. 2024) was observed based on the genotype of SNP markers *SNP-459779133_4A, SNP-459972095_4A*, and *SNP-460562184_4A* taken from DArTSeq genotypic data (Nguyen et al. 2023).

### Pedigree evaluations

The pedigree records of oat lines of interest were obtained from Fitzsimmons et al. (1983), and T3/Oat database (Morales et al. 2022), with information extracted from the “Pedigrees of Oat Lines” POOL database (https://triticeaetoolbox.org/POOL/index_db.php; Tinker and Deyl 2005).

## Results

### Field assessment of rust infection severity

The *Pca* race used in Manjimup fields 2023 and 2024 (MJ23 and MJ24), was 0001-2 and 0005-0, respectively, while the Cobbitty 2024 (CB24) nursery included a mix of races: 0767-3,4,5,6,10,Wa, Vo; 3707-1,4,5,6,7,10,12,Wa,Nu,Gw,Ge,Dr,Al; and 4473-4,6,10, Bett, Barc. The virulence profiles of these races were determined by nursery managers, following the methods described by Park (2000) and presented in **Supplementary File 1 Table S4**. The rust races in CB24 were broadly virulent, exhibiting virulence against 29 differential lines. In contrast, the races in MJ23 and MJ24 are virulent to only Swan, Pc46, and Pc38 in MJ23, and Swan and Pc71 in MJ24.

The RILs from Provena x GS7 and Boyer x GS7 populations and the parental lines were scored for crown rust infection severity (CRS) at the flowering stage in MJ23 and MJ24, and CB24. The data showed rust resistance trait segregation (**Fig. 1**), with a high correlation between crown rust severity percentage in MJ23 and MJ24 for both populations, Provena x GS7 (r = 0.72) and Boyer x GS7 (0.58) (**Supplementary File 1 Table S5).** In contrast, lower correlations were observed between the Manjimup trials (MJ23 and MJ24) and the Cobbitty trial (CB24) (r ≤3.5) (**Supplementary File 1 Table S5**). Rust severity was higher in Cobbitty than in Manjimup, as a greater number of susceptible lines were observed in Cobbitty (**Fig. 1**). Among the parental lines, Provena remained consistently susceptible across trials (CRS = 80–100%), Boyer showed high resistance in all trials (CRS = 0–15%), and GS7 exhibited strong resistance in MJ23 and MJ24 trials (CRS = 0–5%) but displayed a moderate resistance-to-moderate susceptibility in CB24 (CRS = 40%). Across all trials, a greater number of lines with low crown rust severity (CRS = 0 – 40%) were observed in the Boyer x GS7 population, compared to Provena x GS7 (**Fig. 1**), likely reflecting the previously reported combined resistance contribution from both parental lines, Boyer and GS7 (Babiker et al. 2015).

**Fig. 1.**
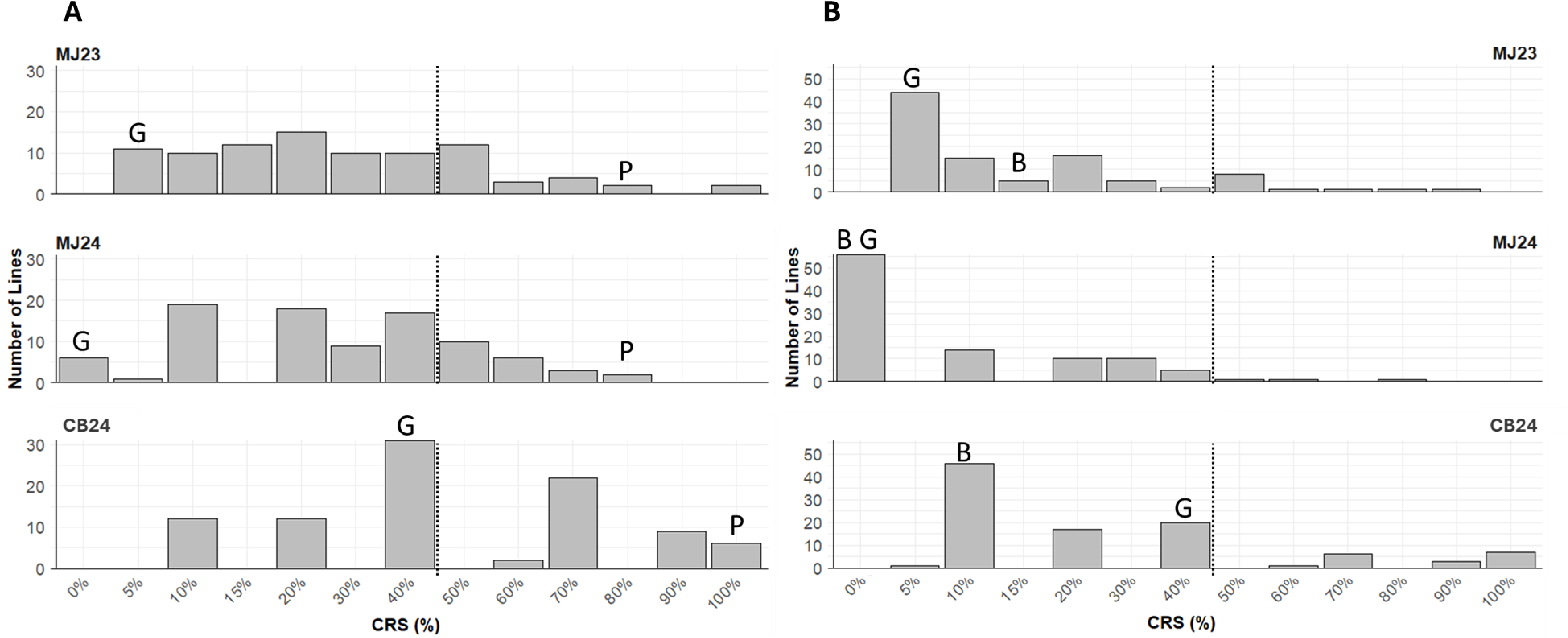
Histogram of field infection severity data from Manjimup 2023 (MJ23), 2024 (MJ24) and Cobbitty 2024 (CB24) of (A) Provena (P) x GS7 (G) RILs (n=91); (B) Boyer (B) x GS7 (G) RILs (n = 98). The x-axis represents crown rust severity = CRS, CRS < 50 % = resistance and CRS ≥ 50 % = susceptibility whereas the y-axis represents number of lines. The dash lines separate resistance (left) and susceptibility (right).

### Construction of genetic maps of Provena x GS7 and Boyer x GS7 RILs

The RILs from Provena x GS7 and Boyer x GS7 mapping populations and the parental lines were genotyped using DArTSeq, generating 31,101 and 26,385 segregating SNPs, respectively. After filtering and converting to a parent-based format (ABH), 5,931 and 6,386 markers were used to construct a genetic map for each family. The final genetic maps included 4,493 and 5,048 SNPs for Provena x GS7 and Boyer x GS7, respectively, with 2,323 common markers between the two populations (**Fig. 2** and **Supplementary File 2**). The genetic maps spanned 2191.5 cM and 2568.2 cM, encompassing 43 and 40 linkage groups with an average marker spacing of 0.5 cM in both maps (**Supplementary File 2** and **Supplementary File 3 Fig. S1**). The DArTSeq markers were also assigned to genomic locations in the *Avena sativa* OT3098 v2 reference genome sequence based on sequence alignment. A high proportion of markers within the same linkage group aligned to the same chromosome on the physical map (OT3098 v2): 96.75% for Provena x GS7 (**Supplementary File 3 Fig. S2**) and 95% for Boyer x GS7 (**Supplementary File 3 Fig. S3**). The average correlation coefficient between genetic and physical map orders for 21 chromosomes was r = 0.86 in both populations. This allowed the genetic maps to be anchored to the 21 oat chromosomes using ASMap R package (Taylor and Butler 2017) and each map provided close to complete coverage of the genome (**Fig. 1**). Chr1D showed the longest linkage group in Provena x GS7 (262.3 cM), while chr2D was the longest in Boyer x GS7 (269.8 cM) (**Fig. 1**).

**Fig. 2.**
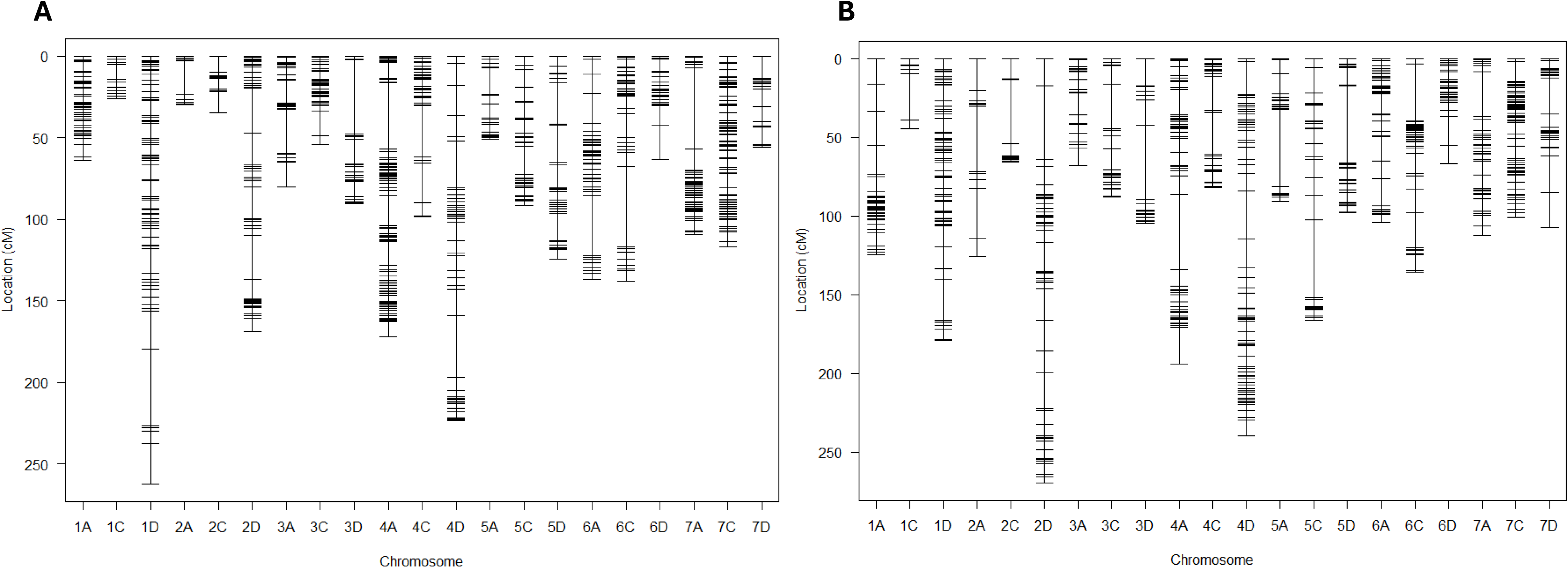
**A** Distribution of markers (4493 SNPs) across 21 oat chromosomes in the genetic map of the Provena x GS7 RIL population (n =91); **B** Distribution of markers (5048 SNPs) across 21 oat chromosomes in the genetic map of the Boyer x GS7 RIL population (n =98). The x-axis represents the 21 oat chromosomes anchored to the genetic maps, while the y-axis shows the genetic positions of markers in centimorgans (cM).

### Identification of QTL associated with oat crown rust severity

#### Provena x GS7 population

QTL analysis identified significant peaks on chr2A (*QPc_Provena_2A*), 4A (*QPc_GS7_4A.1* and *QPc_GS7_4A.2*), 5C (*QPc_GS7_5C*), and 7A (*QPc_GS7_7A* and *QPc_Provena_7A*), with the majority of them carrying resistance alleles from GS7. The exceptions were *QPc_Provena_2A* and *QPc_Provena_7A*, which has the resistance allele from Provena (**Fig. 3** and **Table 1**). *QPc_GS7_4A.2*, located between 158.00 – 162.16 cM, was the only QTL detected in two trials (MJ23 and MJ24) but showed the strongest significance (**Fig. 3A**), accounting for approximately 45% of the genotypic variance explained (GVE) in MJ23 and 35% in MJ24 (**Table 1**). The second most significant detected QTL, *QPc_GS7_4A.1* (LOD = 4.38; GVE = 19.86), which appeared only in the CB24 trial, was located on chr4A between 85.36–104.09 cM, approximately 54 cM away from *QPc_GS7_4A.2*. Low linkage disequilibrium (LD) was found between *QPc_GS7_4A.1* and *QPc_GS7_4A.2* (**Supplementary File 3 Fig. S4A**). The two minor QTL on chr7A, *QPc_GS7_7A* and *QPc_Provena_7A*, were identified in different trials and have low LD to each other (**Supplementary File 3 Fig. S4A**). The *QPc_GS7_7A*, carrying a favourable allele from GS7, was detected in MJ24, whereas *QPc_Provena_7A*, derived from the susceptible parent Provena, was identified in CB24. The two other QTL, *QPc_Provena_2A* and *QPc_GS7_5C*, were both significant in MJ23. *QPc_Provena_2A* was located between 2.89 - 26.4 cM on ch2A, while QPc_GS7_5C was located on chr5C, with GVE values of 14.33% and 16.59%, respectively.

**Fig. 3.**
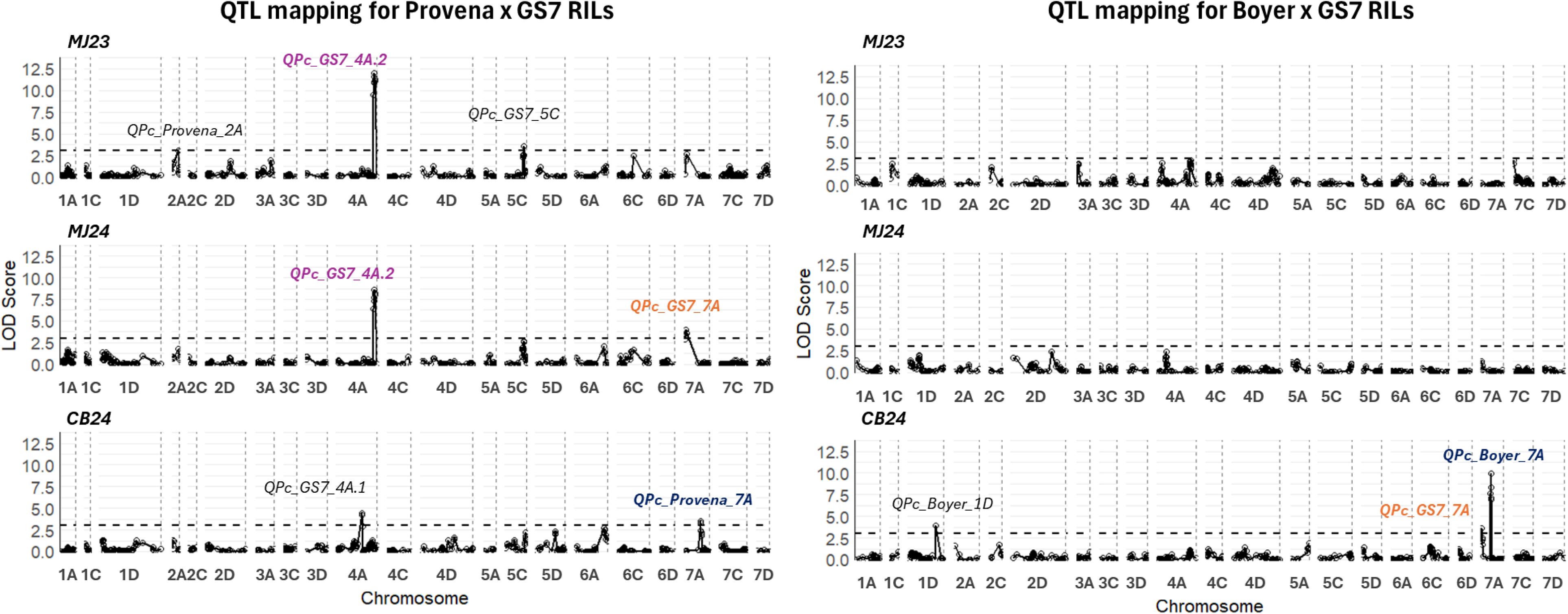
QTL mapping in RIL populations according to field trial. Plots for Provena x GS7 (left) and Boyer x GS7 (right) mapping populations at Manjmup in 2023 (MJ23), 2024 (MJ24), and Cobbitty 2024 (CB24) are shown with chromosome numbers in the x axis and LOD score in the y-axis. LOD threshold (dash lines) = 3. The colours indicate co-located QTL.

**Table 1.** Summary of composite interval mapping in Provena x GS7 and Boyer x GS7 RILs for crown rust severity in Manjmup 2023 (MJ23) - 2024 (MJ24), and Cobbitty 2024 (CB24). Number of * indicates co- located QTL. GVE: Genotypic variance explained.

| Population | QTL | Chromosome | QTL region |  |  | LOD | GVE (%) | Field Trial |
| --- | --- | --- | --- | --- | --- | --- | --- | --- |
|  |  |  | Peak Marker | Peak (cM) | Interval (cM) |  |  |  |
| Provena x GS7 | *QPc_Provena_2A* | 2A | SNP-313607169_2A | 23.46 | 2.89 - 26.4 | 3.06 | 14.33 | MJ23 |
|  | *QPc_GS7_4A.1* | 4A | SNP-352361337_4A | 104.09 | 85.36 - 104.09 | 4.38 | 19.87 | CB24 |
|  | *QPc_GS7_4A.2\** | 4A | SNP-459972095_4A | 161.60 | 158.00 - 162.16 | 11.95-8.18 | 45.36 - 35.23 | MJ23 - MJ24 |
|  | *QPc_GS7_5C* | 5C | SNP-536888142_5C | 80.76 | 75.35 -85.76 | 3.58 | 16.59 | MJ23 |
|  | *QPc_GS7_7A\*\** | 7A | SNP-4039736_7A | 3.60 | 0.00 - 7.03 | 4.01 | 18.38 | MJ24 |
|  | *QPc_Provena_7A\*\*\** | 7A | SNP-66719542_7A | 70.44 | 69.88 - 74.38 | 3.56 | 16.46 | CB24 |
| Boyer x GS7 | *QPc_Boyer_1D* | 1D | SNP-432451680_1D | 140.02 | 133.41 - 166.12 | 3.89 | 16.69 | CB24 |
|  | *QPc_GS7_7A\*\** | 7A | SNP-4039736_7A | 0.64 | 0.00 - 1.92 | 3.57 | 15.46 | CB24 |
|  | *QPc_Boyer_7A\*\*\** | 7A | SNP-66719542_7A | 49.10 | 48.3 - 50.14 | 9.98 | 37.28 | CB24 |

#### Boyer x GS7 populatio

Three significant QTL were identified in the Boyer x GS7 RILs on ch1D (*QPc_Boyer_1D* with GVE = 16.69%) and 7A (*QPc_GS7_7A* with GVE = 37.28% and *QPc_Boyer_7A* with GVE = 15.46%) (**Fig. 3**). All were detected in CB24, while none were identified in MJ23 or MJ24, likely due to the high resistance of most lines, with only a few RILs exhibiting susceptibility in Manjimup trials (**Fig. 1**). The *QPc_GS7_7A* identified in this population, located at 0.00 - 1.92 cM and carrying a favourable allele from GS7, corresponds to the same QTL identified in the Provena x GS7 population (**Table 1**). Similarly, *QPc_Boyer_7A* mapped between 48.3 - 50.14 cM, with a resistance allele from Boyer, co-located with *QPc_Provena_7A*, previously identified in the Provena x GS7 population.

### KASP marker development and screening of the oat collection

Several SNPs that were significantly associated with *QPc_GS7_4A.1*, *QPc_GS7_4A.2*, *QPc_GS7_7A*, and *QPc_Provena_7A*/*QPc_Boyer_7A* were successfully converted into KASP markers, including *KASP_SNP-350765064_4A, KASP_SNP-351486359_4A,* and *KASP_SNP-352361337_4A* for *QPc_GS7_4A.1*, *KASP_SNP-459779133_4A*, *KASP_SNP-459972095_4A*, and *KASP_SNP-460562184_4A* for *QPc_GS7_4A.2*, *KASP_SNP-4039736_7A* and *KASP_SNP-411798_7A* for *QPc_GS7_7A*, and *KASP_SNP-67318388_7A* and *KASP_SNP-69787185_7A* for *QPc_Provena_7A*/*QPc_Boyer_7A* (**Supplementary File 1 Table S3**). KASP assays were performed to assess the presence of these QTL in an oat collection of 182 oat lines (**Supplementary File 1 Table S6**). The results were integrated with rust severity scores of the oat collection from MJ23 and CB24, and a T-test was conducted to evaluate allelic effects on the phenotype (**Fig. 4**).

**Fig. 4.**
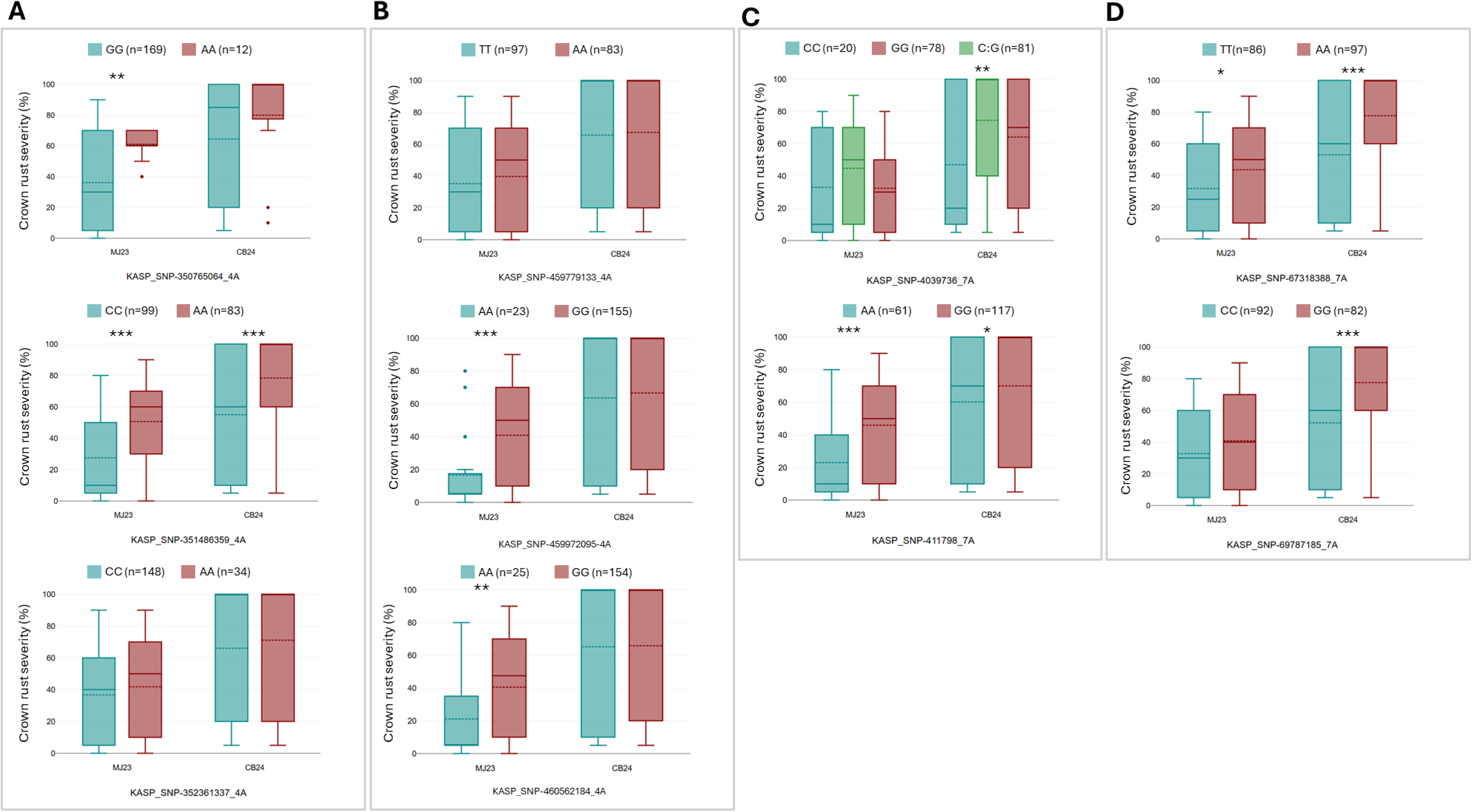
Phenotypic effect of KASPs associated with QTL A QPc_GS7_4A.1; B QPc_GS7_4A.2; C QPc_GS7_7A; D QPc_Provena_7A/QPc_Boyer_7A in the oat collection (n=182). Pairwise t-tests were used to compare the allelic effect on rust severity scores in Manjimup 2023 (MJ23) and Cobbitty 2024 (CB24); Significant levels are based on *p < 0.05, **p < 0.01, ***p < 0.001. The teal colour indicates the resistant alleles, the rose indicates the susceptible alleles and the green indicates the heterozygous allele. The boxes’ solid and dash lines indicate median and mean, respectively).

The markers *KASP_SNP-351486359_4A* (*QPc_GS7_4A.1*), *KASP_SNP-411798_7A* (*QPc_GS7_7A*), and *KASP_SNP-67318388_7A* (*QPc_Provena_7A*/*QPc_Boyer_7A*) showed consistent associations with crown rust severity in the oat collection across both MJ23 and CB24 trials (**Fig. 4**). In contrast, the markers for *QPc_GS7_4A.2*, *KASP_SNP-459972095_4A* and *KASP_SNP-460562184_4A* were only associated with resistance scores in MJ23 but not CB24, aligning with QTL mapping results that identified this QTL only in MJ23 and MJ24.

### QTL *QPc_GS7_4A.2* is colocalised with genomic regions associated with Adult Plant Resistance and other race-specific resistance

To further explore the most significant QTL *QPc_GS7_4A.2* derived from GS7 on chr4A, we examined its colocalization with known resistance loci through comparative mapping analysis. A BLAST search of the flanking sequences of markers significantly associated with rust resistance from previous studies (Babiker et al. 2015; Klos et al. 2017; Nazareno et al. 2022) identified significant matches (% identity > 90.1 and e-value < 2.55E-18) on both chr4A and 4D in reference genome OT3098 v2 (**Supplementary File 1 Table S7** and **Supplementary File 3 Fig. S5**). Additionally, some markers physically mapped to chr4D in our QTL *QPc_GS7_4A.2* overlap with the syntenic APR locus *QPc.APR4D.2* identified by Nazareno et al. (2022) on chr4D (**Supplementary File 3 Fig. S5A**). Synteny analysis revealed significant homology across most regions of chr4A and 4D (**Supplementary File 3 Fig. S5B**). The *QPc_GS7_4A.2* is overlapped with the genomic region associated with the seedling resistance gene *Pc61* and APR loci identified by Klos et al. (2017) and Nazareno et al. (2022), respectively (**Supplementary File 3 Fig. S5A**). Pedigree connections were found between GS7 and both Coker234 (*Pc61*) and Coker227 (*Pc60*) (**Fig 5A**).

**Fig. 5.**
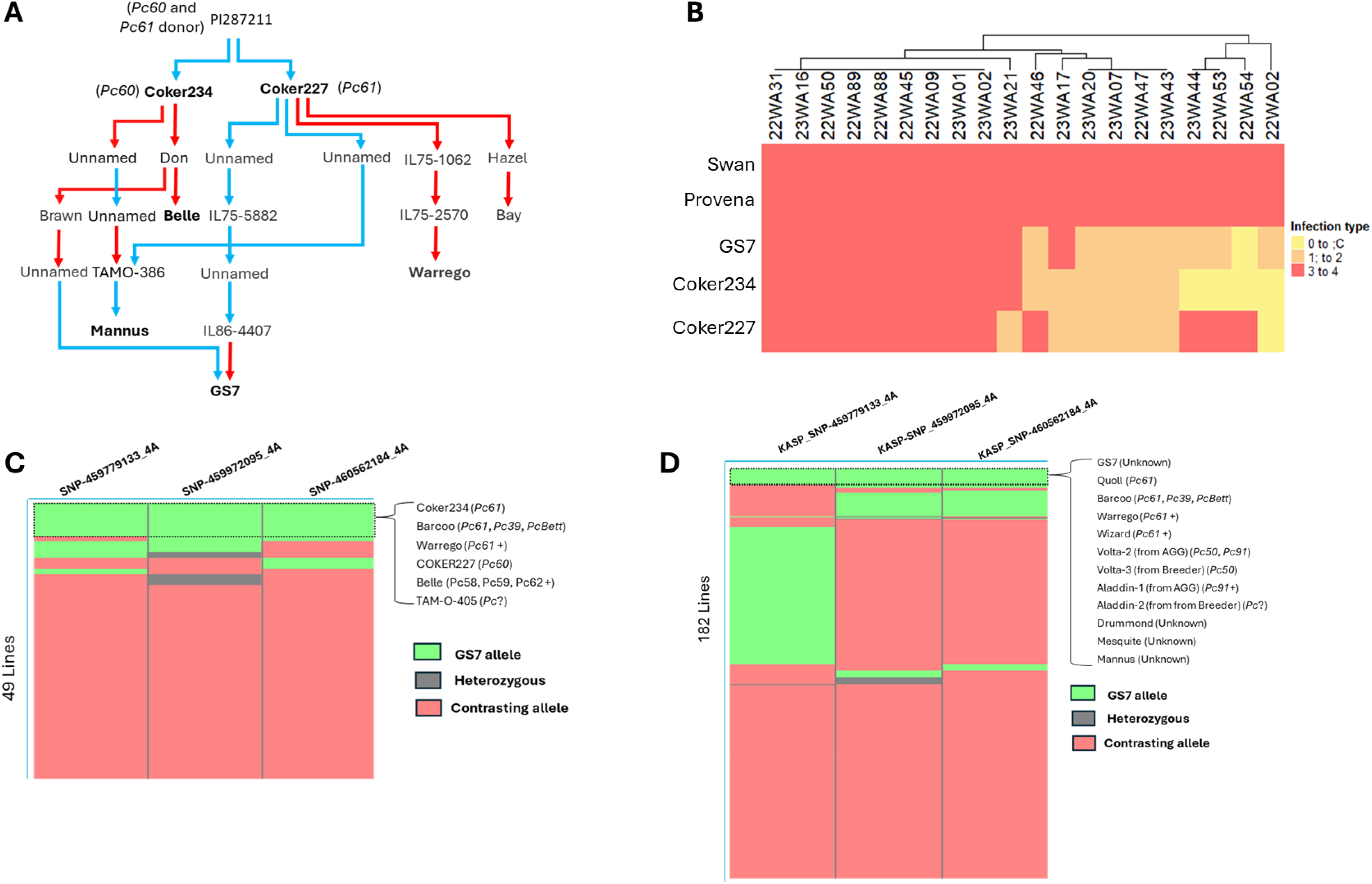
**A** Pedigree relationship of GS7 and Coker234 (Pc61) and Coker227 (Pc60). Pedigrees were obtained from “Pedigrees of Oat Lines” POOL database (https://triticeaetoolbox.org/POOL; Tinker and Deyl, 2005) and Fitzsimmons et al. (1983). Red lines indicate a maternal relationship, and blue indicates a paternal relationship. Name of oat lines in bold represent lines that carry the resistance haplotype at QPc_GS7_4A.2. **B** Heatmap comparing virulence profiles of 20 Pca isolates on GS7, Provena, Pc61, and Pc60. Swan is a widely susceptible cultivar used as the susceptible control. The colour range indicates the infection type of isolate on host: resistant (0 to 2) to susceptible (3 to 4). C Presence of haplotype 4A_GS7 in the oat differential set (n=49) and D the oat collection (n=182). The markers on top are the most significant SNP at the QTL region on chr4A taken from DArTSeq in the differential set, converted to KASP, and tested in the oat collection. Colour indicates the allele of GS7 (green), heterozygous (grey), and the contrasting allele (rose).

Given that *QPc_GS7_4A.2* overlapped with the known *Pc61* locus and manifested only in the Manjimup field across both the Provena x GS7 mapping population and the oat collection, along with GS7’s pedigree connection to both Coker234 (*Pc61*) and Coker227 (*Pc60*), we proposed that this QTL may associate with a seedling resistance gene. A subset of 30 RILs from the Provena x GS7 population, carrying contrasting alleles at the *QPc_GS7_4A.2* locus (15 with the resistance allele and 15 with the susceptible allele), was selected for seedling resistance testing against the rust isolate 22WA54. The results showed that GS7 and the 15 RILs carrying the *QPc_GS7_4A.2* resistance allele exhibited resistance, whereas Provena and the 15 RILs carrying the *QPc_GS7_4A.2* susceptible allele were all susceptible (**Supplementary File 2**). QTL analysis using the seedling resistance data from this subset of samples identified only a single peak that co-mapped with the *QPc_GS7_4A.2* locus (**Supplementary File 3 Fig. S6**), confirming its role as an ASR locus.

Another rust seedling resistance phenotyping experiment at the seedling stage was conducted to compare the resistance profiles of Provena, GS7, Pc60, and Pc61. The experiment used 20 contemporary *Pca* isolates collected from WA in 2022 and 2023 (Henningsen et al. 2024; Nguyen et al. 2025). The result indicated that Provena was susceptible to all tested isolates, while GS7 exhibited seedling resistance to nine *Pca* isolates, confirming it carries at least one ASR gene. Additionally, GS7 displayed a highly similar resistance profile to Coker234 (*Pc61* differential line), differing only for one isolate (**Fig. 5B**).

To confirm the specificity of *QPc_GS7_4A.2* in the oat crown rust differential set (n = 49; **Supplementary File 1 Table S8**), previously used in *Pca* virulence surveillance (Henningsen et al. 2024; Nguyen et al. 2025), we examined the presence of *QPc_GS7_4A.2* resistance haplotype in these lines (**Fig. 5C**). The resistance haplotype of *QPc_GS7_4A.2* are defined by three SNPs previously used to develop KASP assays for *QPc_GS7_4A.2*, *SNP-459779133_4A*, *SNP-459972095_4A*, and *SNP-460562184_4A* that were also found in the DArTSeq data of the oat crown rust differential set (Nguyen et al. 2024). In the differential set, the resistance haplotype of *QPc_GS7_4A.2* was present in six lines, for which the presence of certain race-specific genes has been postulated based on pathogenicity assays (**Fig. 5C**). These lines include Coker234 (*Pc61* differential line), Coker227 (*Pc60* differential line), Warrego (*Pc61*+), Barcoo (*Pc61*, *Pc39*, *PcBett*), Belle (*Pc58*, *Pc59*, *Pc62*, *Pc?*), and TAM-O-405 (Unknown resistance) (Carson 2017; Forsberg et al. 1999; Park et al. 2009). Belle also has a pedigree connection to Coker24 (*Pc61*) (**Fig. 5A**).

In the oat collection, the resistance haplotype of *QPc_GS7_4A.2* was found in 12 lines through genotyping with KASP markers *KASP_SNP-459779133_4A*, *KASP_SNP-459972095_4A*, and *KASP_SNP-460562184_4A* (**Fig. 5D** and **Supplementary File 1 Table S6**). In addition to GS7, eleven other oat lines carried the *4A_GS7* resistance haplotype, including Quoll (*Pc61*), Warrego (*Pc61*+), Barcoo (*Pc61*, *Pc39*, *PcBett*), Wizard (*Pc61*) (Park et al. 2009; Park 2013; Cuddy et al. 2016), Volta-2 (*Pc50*, *Pc91*), Volta-3 (*Pc50*), Aladdin-1 (*Pc91*), Aladdin-2 (unknown *Pc* gene(s)) (Park et al. 2013; Nguyen et al. 2023), Mesquite, Drummond (*Pc39*) (Nguyen et al. 2024), and Mannus. Among these, Mannus was found to have a pedigree connection with Coker234 (*Pc61*) (**Fig. 5A**). Notably, Mesquite is the only postulated APR line in the oat collection that was found to carry the resistance haplotype of *QPc_GS7_4A.2* (**Supplementary File 1 Table S6**). Previously, a QTL in Mesquite was mapped to chr4A by Nazareno et al. (2022), which is 50 Mb from *QPc_GS7_4A.2* (**Supplementary File 3 Fig. S5A**). Two other lines, Amarela and NMO 877, which also had APR QTL mapped to the same region of *QPc_GS7_4A.2* on chr4A (Nazareno et al., 2022 (**Supplementary File 3 Fig. S5A**), differed from the *QPc_GS7_4A.2* resistance haplotype by one marker (**Supplementary File 1 Table S6**).

The *QPc_GS7_4A.2* interval (∼5 Mbp on chr4A, 456,300,687–461,488,017 bp) in the *Avena sativa* OT3098 v2 genome contains 69 annotated genes, including 25 related to disease resistance (**Supplementary File 1 Table S9)**. Notably, a cluster of 10 Disease Resistance Protein RGA5 genes (458,178,696–459,022,508 bp) and a tandem of three ACCELERATED CELL DEATH 6 (ACD6) genes were identified. Other potential candidates include Ankyrin repeat-containing protein NPR4, WRKY transcription factor 49, Receptor-like cytoplasmic kinase 176, and Disease resistance protein Pik-2.

## Discussion

This study employed two established RIL mapping populations (Babika et al. 2015) derived from APR carrying lines, GS7 and Boyer, and a susceptible cultivar (Provena) to assess the effectiveness of the postulated APR loci present in these families under Australian field conditions. Both Boyer and GS7 were confirmed to exhibit high levels of resistance to oat crown rust, while Provena was highly susceptible. QTL analysis using Provena x GS7 and Boyer x GS7 RILs identified multiple loci of interest. Those derived from GS7 were located on chr4A, 5C, and 7A, while loci from Boyer were found on chr1D and 7A, along with two Provena-derived QTL (*QPc_Provena_2A* and *QPc_Provena_7A*). Notably, *QPc_Provena_7A* and *QPc_Boyer_7A* both co-locate with *QCr.cdl11-13A*, a GS7-derived QTL reported by Babiker et al. (2015), which was previously identified in both Provena x GS7 and Boyer x GS7 mapping populations across three trials in the U.S. environment. Another difference between this study and the findings from Babiker et al. (2015) is the detection of *QPc_GS7_4A.1* at 352 Mb in the Provena x GS7 population, a locus previously detected in the Provena × Boyer population in the US, where the resistance allele originated from Boyer. The discrepancy in QTL identification between our study and Babiker et al. (2015) could be attributed to factors such as environmental variation, genotype-environment interactions, QTL epistasis, or differences in pathogen mixtures used at the nurseries during evaluation (Lindhout 2002).

The strongest loci in this study are *QPc_GS7_4A.2*, detected in the trials MJ23 and MJ24, and *QPc_Boyer_7A*, identified in the CB24 trial. The QTL *QPc_GS7_4A.2* overlapped with an APR QTL on chr4A reported by Nazareno et al. (2022) and a QTL region previously associated with the ASR genes *Pc61* and *Pc64* (Klos et al. 2017). Analysis of the *QPc_GS7_4A.2* haplotype at the QTL across the differential set identified the Pc61 differential line (Coker234) as one of the carriers, but not the Pc64 differential line. In the oat collection, the *QPc_GS7_4A.2* resistance haplotype was also present in other lines postulated to carry *Pc61*, such as Quoll, Wizard, Warrego, and Barcoo. The rust phenotyping experiment at the seedling stage using 20 *Pca* isolates showed similar resistance profiles between GS7 and the Pc61 differential line. The seedling resistance assay and subsequent QTL mapping analysis using 30 RILs carrying contrasting alleles at *QPc_GS7_4A.2* further confirmed the association of this locus with seedling resistance. All these findings, along with Klos et al. (2017) identifying the same genomic region linked to *Pc61*, strongly suggest that the mapped QTL in GS7 is closely linked to the *Pc61* locus or potentially represents *Pc61* itself. Furthermore, based on the distribution of virulent isolates across trials, *QPc_GS7_4A.2* was effective in MJ23 and MJ24, but not in CB24, likely due to the presence of *Pc61*-virulent pathotypes in CB24, which were absent in the MJ23 trial.

In CB24, the resistance in the Provena x GS7 RILs was attributed to the QTL *QPc_GS7_4A.1* and *QPc_Provena_7A*. In the oat collection, the *QPc_GS7_4A.1* locus was significantly associated with resistance in both MJ23 and CB24, suggesting it may be a potential APR locus effective across multiple environments and diverse Pca isolates. This locus is collocated with previously identified APR loci identified by Babiker et al. (2015) and Nazareno et al. (2022). The findings *QPc_GS7_4A.1* and *QPc_GS7_4A.2* in GS7 and their difference in effectiveness across environments indicate that GS7 carries both ASR (*Pc61*) and APR. The co-presence of ASR gene with APR loci has been previously reported. For example, the oat line Garry carries multiple ASR genes (*Pc24*, *Pc25*, *Pc26*) alongside APR genes (*Pc27* and *Pc28*), and the oat line TAM O-301 harbors *Pc58* and additional APR-contributing genes (Upadhyaya and Baker 1960, 1962; Cason 2017; Hoffman et al. 2006). Co-location of APR and seedling resistance genes has also been documented in wheat. For example, *Lr12* (APR) and *Lr31* (ASR) both map to chromosome 4B, and their complementary interaction contributes to resistance (Singh et al. 1999). Similarly, a major field stem rust resistance gene co-locates with *Sr12* in ‘Thatcher’ wheat (Hiebert et al. 2016).

The oat line Coker227 (*Pc60*), which shares a high genetic similarity with Coker234 (*Pc61*) (Nguyen et al. 2024) but showed a different seedling resistance profile, also carries the *QPc_GS7_4A.2* resistance haplotype. This is likely a result of GS7’s descent from Pc60 in the pedigree, with both Coker227 (*Pc60*) and Coker234 (*Pc61)* originating from a common source *A*. *sterilis* PI 287211 (Carson 2017). Moreover, the *QPc_GS7_4A.2* resistance haplotype was also found in several crown rust-resistant lines not yet catalogued to carry any known *Pc* gene TAM-O-405, Drummond, and Mannus. Of these three lines, Mannus has pedigree connection with Coker234 (*Pc61)*. These lines may potentially harbor the *QPc_GS7_4A.2* as part of their resistance mechanism, though further studies are needed to confirm this. Oat cultivars Volta-2 and Aladdin-1, previously postulated to carry *Pc91* (Nguyen et al. 2023), were also found to harbor the *QPc_GS7_4A.2* resistance haplotype. Interestingly, a recent genome-wide association study identified a shared genomic interval in the *Pca* genome significantly associated with virulence to both *Pc61* and *Pc91* (Hewitt et al. 2024), suggesting a possible mechanistic link between these resistance loci in oats. Our findings align with this observation, supporting a connection between *Pc61* and *Pc91*.

A candidate gene search identified 25 of 69 annotated genes in the 4A QTL interval that are putatively linked to disease resistance mechanisms. Notably, a cluster of 10 Disease Resistance RGA5 and three *ACD6* genes on chromosome 4A were found in this region. These RGA5 belong to the gene family that encodes NB-LRR proteins in rice, known to mediate resistance to the fungal pathogen *Magnaporthe oryzae,* and exhibit diverse functions in *Avr* recognition (Césari et al. 2014). On the other hand, in Arabidopsis, an *ACD6* was recently identified as the causal gene for leaf senescence (Jasinski et al. 2021). Notably, a gene annotated as Pik2 was identified approximately 2 Mb from the RGA5 cluster within the QTL interval. Pik2, like RGA5, is an NLR protein, and they are functionally related. In rice, RGA5 acts as the sensor in the sensor-helper pair RGA5-RGA4, while Pik2 serves as the helper in the Pik1-Pik2 pair (Zdrzałek et al. 2020). These genetically linked NLR pairs typically operate as sensor-helper systems, where one component recognises pathogen effectors and the other activates immune responses (Zhai et al. 2011; Zdrzałek et al. 2020). In addition, a putative protein WRK49 was also found. In wheat, WRK49 was claimed to confer differential high-temperature seedling-plant resistance to *Puccinia striiformis* f. sp. *tritici* (*Pst)* (Wang et al. 2017). Furthermore, there is a putative protein Ankyrin repeat-containing protein NPR4 in the QLT region, which was orthologous to *TaANKTM1B-4, TaANKTM2B-1, TaANKTM1D-6, TaANKTM3B-9,* TaANKTM4B-5 in wheat. The ankyrin-transmembrane (ANKTM) subfamily is the most abundant subgroup of the ANK superfamily, with roles in pathogen defense (Hu et al. 2022). A homolog of these genes, *TaANKTM2A-5* was found to regulate powdery mildew resistance in wheat (Hu et al. 2022). The candidate genes listed here could be potential targets for cloning from the resistant parent to support functional characterisation.

In conclusion, this study emphasises the potential of GS7 and Boyer as a useful source of crown rust resistance in Australia. The QTL *QPc_GS7_4A.*2 is a seedling resistance locus that is closely linked to *Pc61* and potentially *Pc61* itself. The KASP markers were developed for *QPc_GS7_4A.1, QPc_GS7_4A.2, QPc_GS7_7A,* and *QPc_Provena_7A/QPc_Boyer_7A* will be valuable for marker-assisted breeding in oat improvement programs. Future research should focus on stacking these QTL to evaluate their interaction in a common background, while fine mapping is needed to pinpoint their genetic mechanism.

## Statements & Declarations

## Acknowledgments

We acknowledge the Australian Grains Genebank, Shahryar F. Kianian and Tyler Gordon at US Department of Agriculture-Agricultural Research Service for research discussions, and the GRDC-funded oat phenology project team (grant CSP2007) for contributing oat germplasm.

## Author contributions

This study was planned and designed by MF and PND. AR conducted rust infection phenotyping in the field. DTN performed the genotypic analysis. DL and ECH implemented virulence assessments for the plants at the seedling stage in the growth cabinets. RM assisted with marker design. JS supported computational analyses. DTN, PND and MF wrote the first draft of the manuscript. All authors contributed to the interpretation of results, reviewed the manuscript, and approved the final version.

## Data availability

The DArTSeq genotypes of the oat lines included in this study are deposited in the CSIRO Data Access Portal repository https://doi.org/10.25919/x7sk-qp24. The scripts for genotypic data analysis and QTL mapping are available on GitHub at https://github.com/duongnguyen1987/QTL_mapping.

## Literature Cited

Altschul SF, Gish W, Miller W, Myers EW, Lipman DJ (1990) Basic local alignment search tool. J. Mol. Biol. 215:403–410. 10.1016/S0022-2836(05)80360-2

Ao Y, Li Z, Feng D, et al (2014) *OsCERK1* and *OsRLCK176* play important roles in peptidoglycan and chitin signaling in rice innate immunity. Plant J 80:1072–1084. 10.1111/tpj.12710

Babiker EM, Gordon TC, Jackson EW, Chao S, Harrison SA, Carson ML, Obert DE., Bonman, JM (2015). Quantitative trait loci from two genotypes of oat (*Avena sativa*) conditioning resistance to *Puccinia coronata*. Phytopathology 105, 239–245. 10.1094/PHYTO-04-14-0114-R

Bariana HS, Miah H, Brown GN, Willey N, Lehmensiek A (2007) Molecular mapping of durable rust resistance in wheat and its implication in dreeding. In: Buck, H.T., Nisi, J.E., Salomón, N. (eds) Wheat production in stressed environments. Developments in Plant Breeding, vol 12. Springer, Dordrecht. 10.1007/1-4020-5497-1_88

Broman KW, Wu H, Sen Ś, Churchill GA (2003) R/qtl: QTL mapping in experimental crosses. Bioinformatics 19:889–890. 10.1093/bioinformatics/btg112

Broman KW, Sen S (2009). A Guide to QTL Mapping with R/qtl. Springer, New York. 10.1007/978-0-387-92125-9

Bradbury PJ, Zhang Z, Kroon DE, Casstevens TM, Ramdoss Y, Buckler ES (2007) TASSEL: software for association mapping of complex traits in diverse samples. Bioinformatics 23:2633–2635. 10.1093/bioinformatics/btm308

Cabral AL, Singh D, Park RF (2011) Identification and genetic characterisation of adult plant resistance to crown rust in diploid and tetraploid accessions of *Avena*. Annal Appl Biol 159:220–228. 10.1111/j.1744-7348.2011.00492.x

Carson ML (2008) Virulence frequencies in oat crown fust in the United States from 2001 through 2005. Plant Dis 92:379–384. 10.1094/PDIS-92-3-0379

Carson ML (2017) Oat crown rust. Cereal Disease Lab, Agricultural Research Service, USDA. https://www.ars.usda.gov/midwest-area/stpaul/cereal-disease-lab/docs/cereal-rusts/oat-crown-rust/

Césari S, Kanzaki H, Fujiwara T, et al (2014) The NB-LRR proteins RGA4 and RGA5 interact functionally and physically to confer disease resistance. EMBO J. 33:1941–1959. 10.15252/embj.201487923

Chaffin AS, Huang YF, Smith S, et al (2016) A consensus map in cultivated hexaploid oat reveals conserved grass synteny with substantial subgenome rearrangement. Plant genome 9:10.3835. 10.3835/plantgenome2015.10.0102

Cuddy W, Park R, Bariana H, Bansal U, Singh D, Roake J, Platz G (2016) Cereal rust report 2016. Plant Breeding Institute, University of Sydney.

Dodds PN, Chen J, Outram MA (2024). Pathogen perception and signaling in plant immunity. The Plant cell 36:1465–1481. 10.1093/plcell/koae020

Dodds PN (2023) From gene-for-gene to resistosomes: Flor’s Enduring Legacy. Molecular Plant-Microbe Interactions 36:461–446. 10.1094/MPMI-06-23-0081-HH

Ellis JG, Lagudah ES, Spielmeyer W, Dodds PN (2014). The past, present and future of breeding for rust resistance in wheat. Frontiers in Plant Science 5: 641. 10.3389/fpls.2014.00641

Forsberg RA; Kaeppler HF, Duerst RD (1999) Registration of Belle oat. Crop Science 39:878–879. 10.2135/cropsci1999.0011183X003900030058x

Figueroa M, Dodds PN, Henningsen EC (2020) Evolution of virulence in rust fungi — multiple solutions to one problem. Curr. Opin. Plant Biol 56: 20–27. 10.1016/j.pbi.2020.02.007

Fitzsimmons RW, Roberts GL., Wrigley CW (1983). Australian oat varieties (Melbourne: CSIRO Publishing). 10.1071/9780643105447

Furuta T, Ashikari M, Jena KK, Doi K, Reuscher S. Adapting genotyping-by-sequencing for rice F2 populations (2017) G3 (Bethesda) 7:881-893. 10.1534/g3.116.038190

Gruber B, Unmack PJ, Berry OF, Georges A (2018) dartr: An r package to facilitate analysis of SNP data generated from reduced representation genome sequencing. Mol Ecol Resour 18: 691–699. 10.1111/1755-0998.12745

Gu Z (2022) Complex heatmap visualization. IMeta. 1(3). 10.1002/imt2.43

Harder DE, McKenzie RIH, Martens JW (1984) Inheritance of adult plant resistance to crown rust in an accession of *Avena sterilis*. Phytopathology, 74:352–353. 10.1094/Phyto-74-352

Haug-Baltzell A, Stephens SA, Davey S, Scheidegger CE, Lyons E (2017) SynMap2 and SynMap3D: web-based whole-genome synteny browsers. Bioinformatics 33: 2197–2198. 10.1093/bioinformatics/btx144

Heagle AS and Moore MB (1970) Some effects of moderate adult resistance to crown rust of oats. Phytopathology 60:461–466. 10.1094/Phyto-60-461

Henningsen EC, Lewis DC, Nguyen DT et al (2024) Virulence patterns of oat crown rust in Australia - season 2022. Plant Dis 108:1959–1963. 10.1094/PDIS-09-23-1973-SC

Hewitt T, Henningsen EC, Pereira DA et al (2024) Genome-enabled analysis of population dynamics and virulence associated loci in the oat crown rust fungus *Puccinia coronata* f. sp. *avenae*. MPMI. 10.1094/mpmi-09-23-0126-fi

Hiebert CW, Kolmer JA, McCartney CA, Briggs J, Fetch T, Bariana H, Choulet F, Rouse MN, Spielmeyer W (2016). Major gene for field stem rust resistance co-locates with resistance gene *Sr12* in ‘Thatcher’ Wheat. PLoS One. 11:e0157029. 10.1371/journal.pone.0157029

Hu P, Ren Y, Xu J, Wei Q, Song P, Guan Y, Gao H, Zhang Y, Hu H and Li C (2022) Identification of ankyrin-transmembrane-type subfamily genes in *Triticeae* species reveals *TaANKTM2A-5* regulates powdery mildew resistance in wheat. Front. Plant Sci. 13:943217. 10.3389/fpls.2022.943217

Hoffman DL, Chong J, Jackson EW, Obert DE (2006) Characterization and mapping of a crown rust resistance gene complex (*Pc58*) in TAM O-301. Crop Sci. 46:2630–2635. 10.2135/cropsci2006.01.0014

Jackson EW, Obert DE, Menz M, et al (2007) Characterization and mapping of oat crown rust resistance genes using three assessment methods. Phytopathology 97:1063–1070. 10.1094/PHYTO-97-9-1063

Jasinski S, Fabrissin I, Masson A, Marmagne A, Lécureuil A, Bill L and Chardon F (2021) ACCELERATED CELL DEATH 6 acts on natural leaf senescence and nitrogen fluxes in *Arabidopsis*. Front. Plant Sci. 11:611170. 10.3389/fpls.2020.611170

Jellen EN, Wight CP, Spannagl M, Blake VC, Chong J, Herrmann MH, Howarth CJ, Huang Y-F, Juqing J, Katsiotis A, Langdon T, Li C, Park R, Tinker NA, Sen TZ (2024) A uniform gene and chromosome nomenclature system for oat (*Avena* ssp.) Crop and Pasture Science. 10.1071/CP23247

Jones JDG, Staskawicz, BJ, Dangl, JL (2024) The plant immune system: From discovery to deployment. Cell, 187: 2095–2116. 10.1016/j.cell.2024.03.045

Klos KE, Yimer BA, Babiker EM, Beattie AD, Bonman JM, Carson ML et al (2017) Genome-wide association mapping of crown rust resistance in oat elite germplasm. The Plant Genome, 10(2), 10.3835/plantgenome2016.10.0107

Krattinger SG, Lagudah ES, Spielmeyer W, Singh RP, Huerta-Espino J, McFadden H et al (2009) A putative ABC transporter confers durable resistance to multiple fungal pathogens in wheat. Science 323:1360–1363 10.1126/science.1166453

Krattinger SG, Sucher J, Selter LL, Chauhan H, Zhou B, Tang M et al (2016) The wheat durable, multipathogen resistance gene *Lr34* confers partial blast resistance in rice. Plant Biotechnol J 14:1261–1268 10.1111/pbi.12491

Lin Y, Gnanesh BN, Chong J et al (2014) A major quantitative trait locus conferring adult plant partial resistance to crown rust in oat. BMC Plant Biol 14, 250. 10.1186/s12870-014-0250-2

Lindhout P (2002) The perspectives of polygenic resistance in breeding for durable disease resistance. Euphytica 124:217–226. 10.1023/A:1015686601404

Luke HH, Chapman WH, Barnett RD (1972) Horizontal resistance of Red Rustproof oats to crown rust. Phytopathology 62:414–417. 10.1094/Phyto-62-414

Miller ME, Nazareno ES, Rottschaefer SM, Riddle J, Dos Santos Pereira D, Li F, … Figueroa M (2020). Increased virulence of *Puccinia coronata* f. sp. *avenae* populations through allele frequency changes at multiple putative *Avr* loci. PLoS Genetics, 16(12), e1009291. 10.1371/journal.pgen.1009291

Milne I, Shaw P, Stephen G, Bayer M, Cardle L, Thomas WTB, Flavell AJ and Marshall D (2010) Flapjack – graphical genotype visualization. Bioinformatics 26: 3133–3134. 10.1093/bioinformatics/btq580

Moore JW, Herrera-Foessel S, Lan C, Schnippenkoetter W, Ayliffe M, Espino-Huerta J et al (2015) A recently evolved hexose transporter variant confers resistance to multiple pathogens in wheat. Nat. Genet. 47:1494–1498. 10.1038/ng.3439

Morales N, Ogbonna AC, Ellerbrock BJ, Bauchet GJ, Tantikanjana T,… Mueller LA (2022) Breedbase: a digital ecosystem for modern plant breeding. G3 (Bethesda, Md.), 12(7), jkac078. 10.1093/g3journal/jkac078

Milne RJ, Dibley KE, Schnippenkoetter W, Mascher M, Lui ACW, Wang L et al (2019) The wheat *Lr67* gene from the sugar transport protein 13 family confers multipathogen resistance in barley. Plant Physiol. 179, 1285–1297. 10.1104/pp.18.00945

Nazareno ES, Li F, Smith M et al (2018) *Puccinia coronata* f. sp. *avenae*: a threat to global oat production. Mol Plant Pathol 19:1047–1060. 10.1111/mpp.12608

Nazareno ES, Fiedler JD, Miller ME et al (2022) A reference-anchored oat linkage map reveals quantitative trait loci conferring adult plant resistance to crown rust (*Puccinia coronata* f. sp. *avenae*). Theor Appl Genet 135:3307–3321. 10.1007/s00122-022-04128-6

Nguyen DT, Henningsen EC, Lewis D et al (2024) Genotypic and resistance profile analysis of two oat crown rust differential sets urges coordination and standardisation. Phytopathology 114:1356–1365. 10.1094/phyto-10-23-0353-r

Nguyen DT, Henningsen EC, Lewis D, Mago R, Sperschneider J, Stone E, Dodds PN, Figueroa M (2025) Characterisation of virulence of Puccinia coronata f. sp. avenae in Australia in the 2023 growing season. Plant Dis. Online ahead of print. 10.1094/PDIS-09-24-2049-SC

Nguyen DT, Lewis D, Henningsen EC, Mago R, Sperschneider J, Dodds PN, Figueroa M (2024) Identification of diagnostic KASP markers for selection of crown rust resistance in oats. bioRxiv 2024.10.27.620528. 10.1101/2024.10.27.620528

Park R (2000). Occurrence and Pathogenic Specialisation in Puccinia coronata in Australasia, 1999-2000. Oat Newsletter. 46. https://wheat.pw.usda.gov/ggpages/oatnewsletter/v46/#Park

Park R, Wellings C, Bariana H, Bansai U (2009) Australia cereal cultivar pedigree and seedling rust genotype information. Cereal Rust Report 2009. Plant Breeding Institute, University of Sydney. https://www.sydney.edu.au/content/dam/corporate/documents/sydney-institute-of-agriculture/research/plant-breeding-and-production/cereal_rust_report_2009_vol_7_2.pdf

Park R (2013) New oat crown rust pathotype with virulence for *Pc91*. Cereal Rust Report 2013. Plant Breeding Institute, University of Sydney. https://www.sydney.edu.au/content/dam/corporate/documents/sydney-institute-of-agriculture/research/plant-breeding-and-production/cereal_rust_report_2013_vol_11_1.pdf

Peterson RF, Campbell AB, and Hannah AE. (1948) A diagrammatic scale for estimating rust intensity on leaves and stems of cereals. Can J Res Sect 26: 496–500. 10.1139/cjr48c-033

Periyannan S, Milne RJ, Figueroa M, Lagudah ES, Dodds PN (2017). An overview of genetic rust resistance: From broad to specific mechanisms. PLOS Pathog. 13(7), e1006380. 10.1371/journal.ppat.1006380

Portyanko VA., Hoffman DL, Lee M, Holland JB (2001) A linkage map of hexaploid oat based on grass anchor DNA clones and its relationship to other oat maps. Genome, 44:249–265. 10.1139/g01-003

Portyanko VA, Chen G, Rines HW, Phillips RL, Leonard KJ, Ochocki GE, Stuthman DD (2005) Quantitative trait loci for partial resistance to crown rust, *Puccinia coronata*, in cultivated oat, *Avena sativa* L. Theor Appl Genet 111:313–324. 10.1007/s00122-005-2024-6

Rines HW, Miller ME, Carson M, Chao S, Tiede T, Wiersma J, Kianian SF (2018) Identification, introgression, and molecular marker genetic analysis and selection of a highly effective novel oat crown rust resistance from diploid oat, *Avena strigosa*. Theor Appl Genet 131:721–733.

Sanz MJ, Jelle, EN, Loarce Y et al (2010) A new chromosome nomenclature system for oat (*Avena sativa* L. and *A. byzantina* C. Koch) based on FISH analysis of monosomic lines. Theor Appl Genet 121: 1541–1552 10.1007/s00122-010-1409-3

Shi P, Shen X, Chen J-C et al (2023) KASP genotyping and semi-quantitation of G275E mutation in the α6 subunit of Thrips palminAChR gene conferring spinetoram resistance. Pest Manag Sci 79:1777–1782. 10.1002/ps.7353

Singh D, Park R, McIntosh RA (1999) Genetic relationship between the adult plant resistance gene *Lr12* and the complementary gene *Lr31* for seedling resistance to leaf rust in common wheat. Plant Pathology: 48:567 –573. 10.1046/j.1365-3059.1999.00391.x

Simons MD (1985) Crown rust. Elsevier eBooks 131–172. 10.1016/b978-0-12-148402-6.50013-4

Taylor J, Butler D (2017) R Package ASMap: Efficient Genetic Linkage Map Construction and Diagnosis. Journal of Statistical Software, 79(6), 1–29. 10.18637/jss.v079.i06

Tinker NA, Deyl JK. (2005) A curated internet database of oat pedigrees. Crop Sci. 45: 2269–2272. 10.2135/cropsci2004.0687

Upadhyaya YM, Baker EP (1960) Studies on the mode of inheritance of Hajira type stem rust resistance and Victoria type crown rust resistance as exhibited in crosses involving the oat variety Garry. Proc Linnean Soc NSW 85:157–179

Upadhyaya YM, Baker EP (1962) Studies on the inheritance of rust resistance in oats. IE The mode of inheritance of crown rust resistance in the varieties Landhafer, Santa Fe, Mutica Ukraine, Trispernia, and Victoria in their crosses with susceptible varieties. Proc Linnean Soc NSW 87:200–219

Wang J, Tao F, Tian W, et al (2017) The wheat WRKY transcription factors TaWRKY49 and TaWRKY62 confer differential high-temperature seedling-plant resistance to Puccinia striiformis f. sp. tritici. PLoS One 12:e0181963. 10.1371/journal.pone.0181963

Welsh JN, Carson RB, Cherewick WJ, Hagborg WAF, Peturson B, Wallace HAH (1953). Oat varieties-past and present. Canada Department of Agriculture, Ottawa, Pub 891

Wickham H (2016). ggplot2: Elegant Graphics for Data Analysis. Springer-Verlag New York. ISBN 978-3-319-24277-4, https://ggplot2.tidyverse.org.

Yao E, Blake VC, Cooper L, Wight CP, Michel S, Cagirici HB, Lazo GR, Birkett CL, Waring DJ, Jannink JL, Holmes I, Waters AJ, Eickholt DP, Sen TZ (2022) GrainGenes: a data-rich repository for small grains genetics and genomics. Database: the journal of biological databases and curation, 2022, baac034. 10.1093/database/baac034

Zadoks JC, Chang TT, Konzak CF (1974) A decimal growth code for the growth stages of cereals. Weed Res 14:415–421. 10.1111/j.1365-3180.1974.tb01084.x

Zhai C, Lin F, Dong Z, et al (2011) The isolation and characterization of Pik, a rice blast resistance gene which emerged after rice domestication. New Phytol 189:321–334. 10.1111/j.1469-8137.2010.03462.x

Zdrzałek R, Kamoun S, Terauchi R, Saitoh H, Banfield MJ (2020) The rice NLR pair *Pikp-1*/*Pikp-2* initiates cell death through receptor cooperation rather than negative regulation. PLoS One 15:e0238616. 10.1371/journal.pone.0238616

