## Supplementary figures and images for "QTL mapping of oat crown rust resistance in Australian fields and identification of a seedling resistance locus in oat line GS7"

### Supplemental Figures

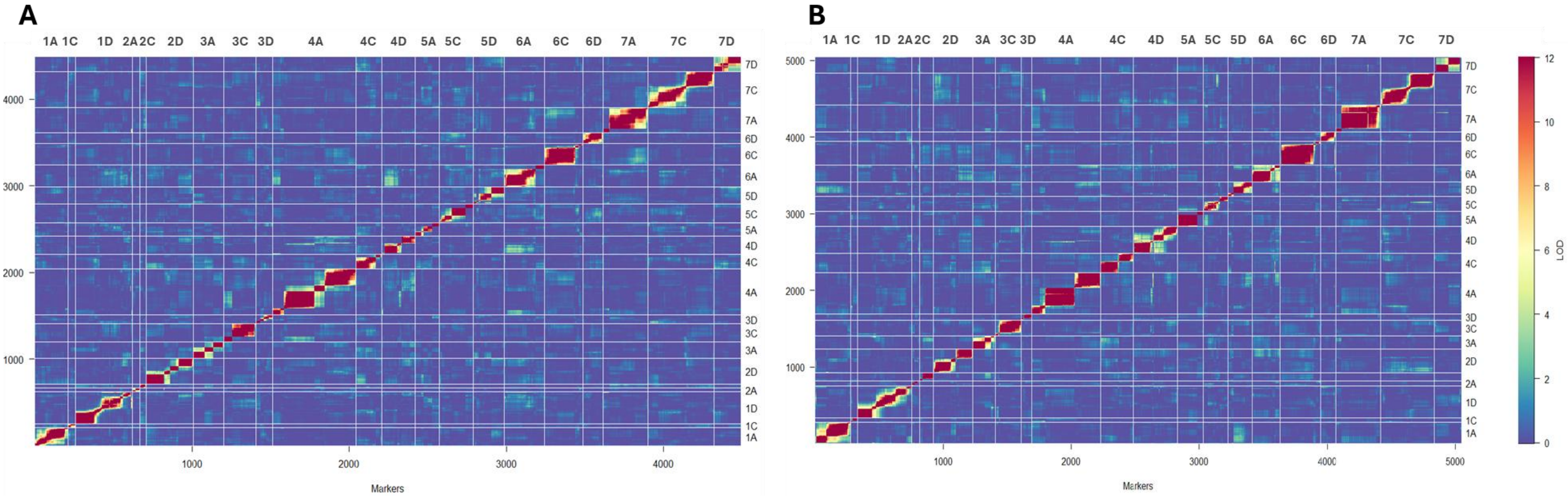

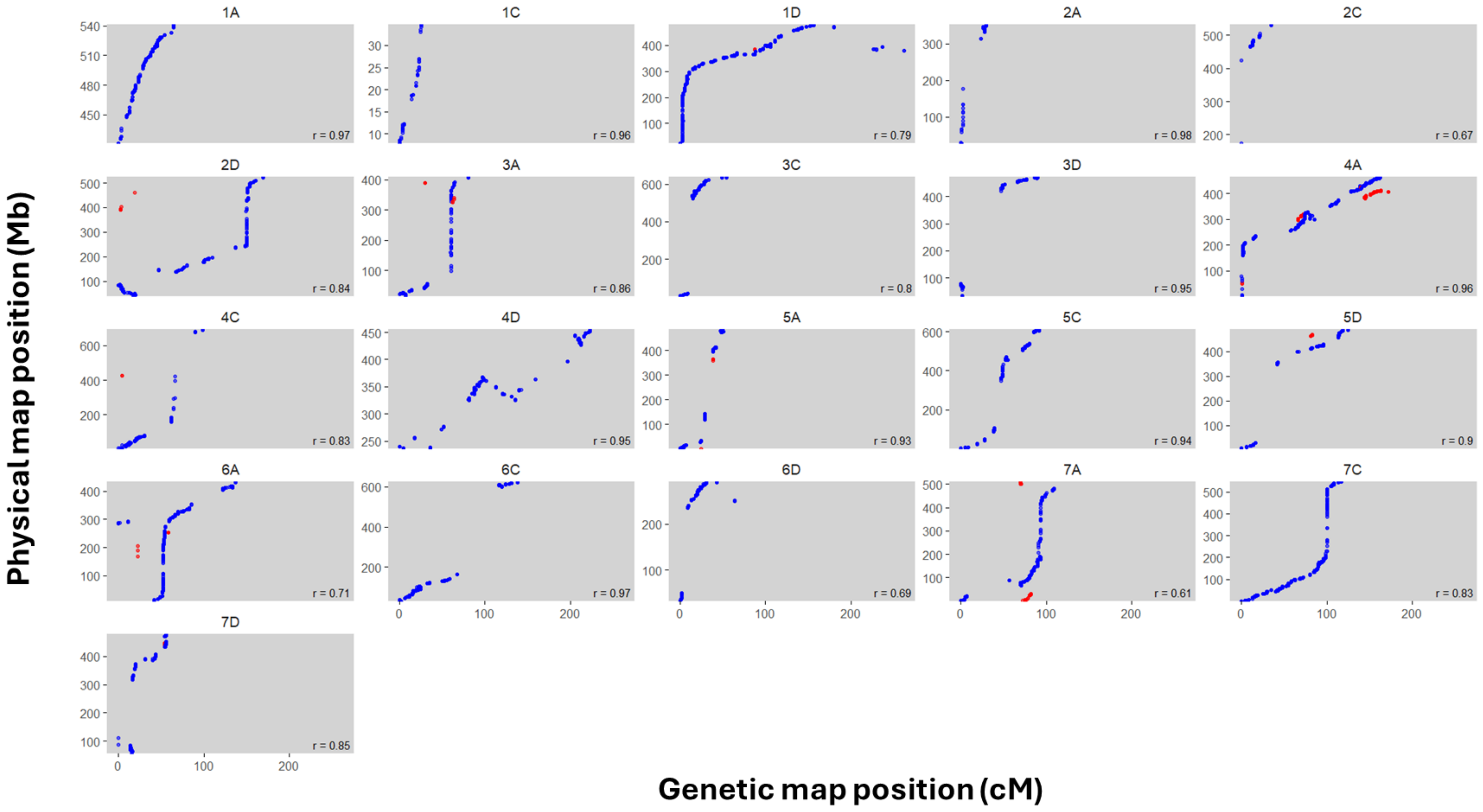

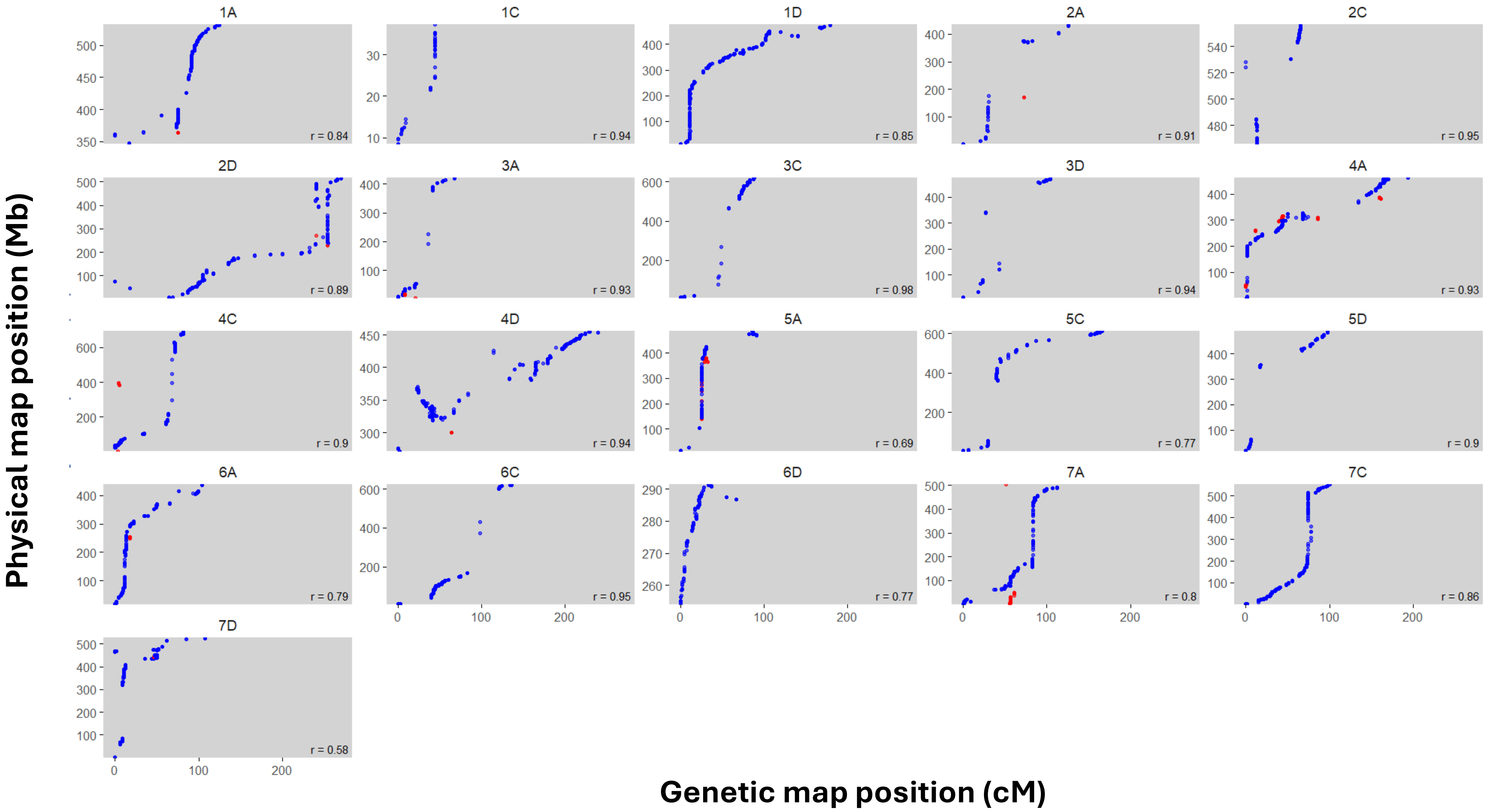

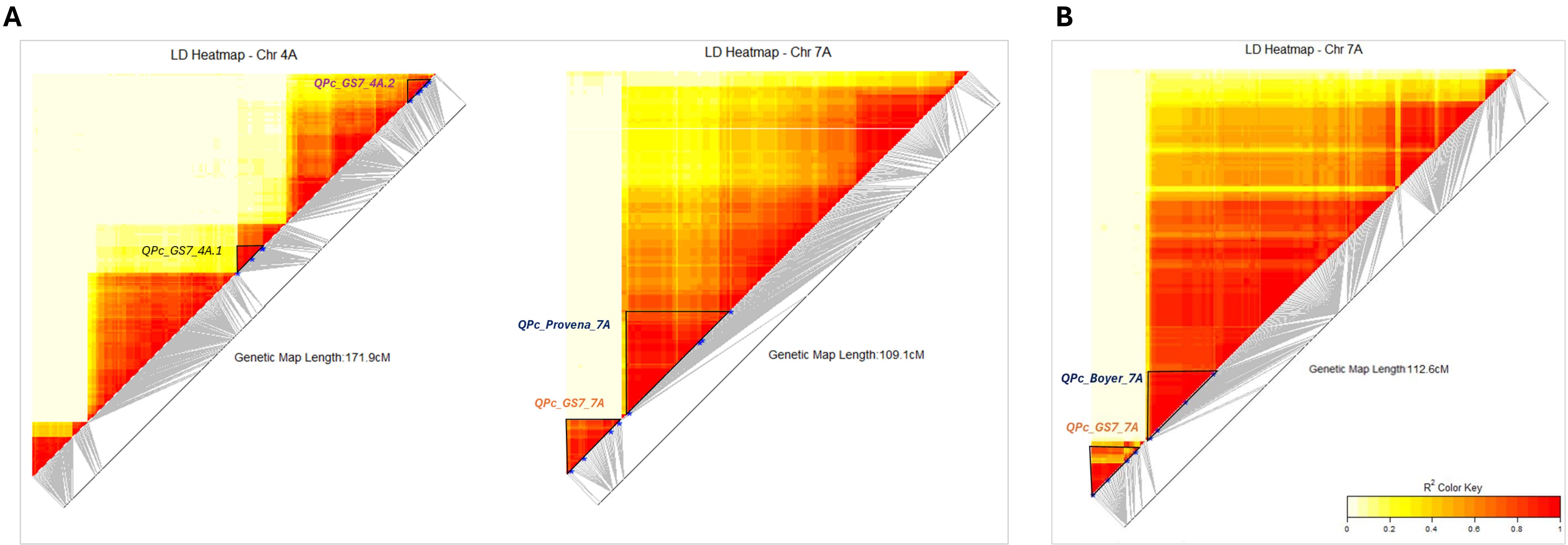

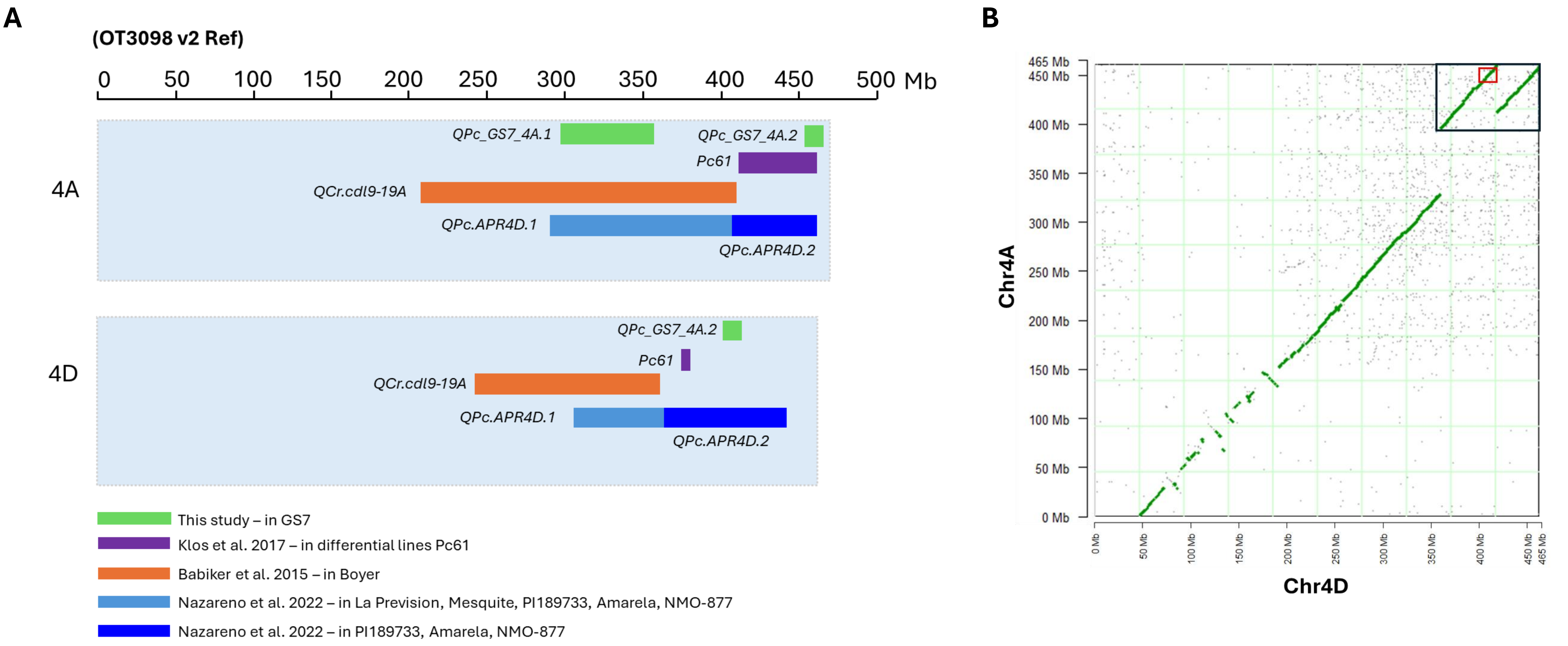

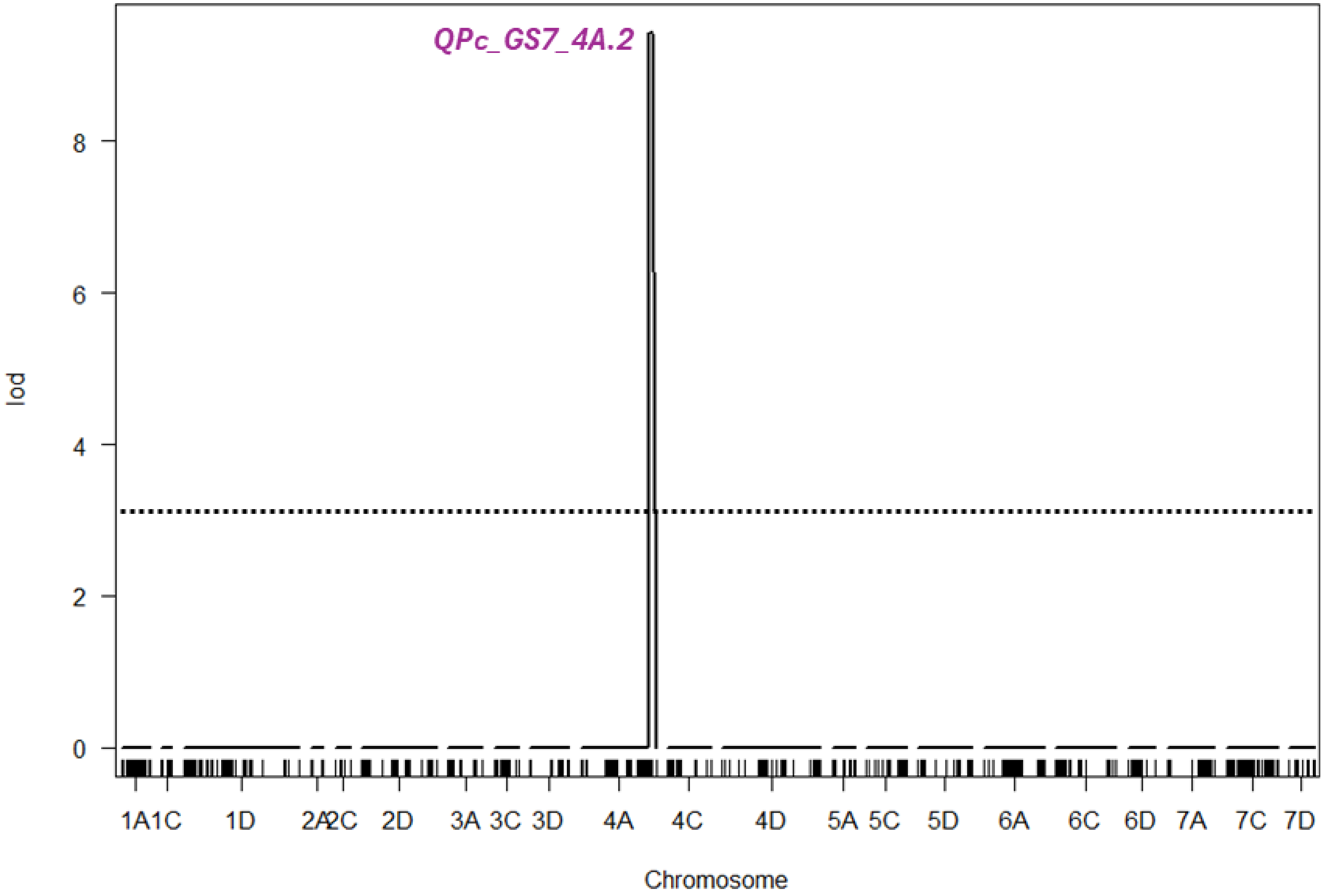
